# Population genomics insights into the recent evolution of SARS-CoV-2

**DOI:** 10.1101/2020.04.21.054122

**Authors:** Maria Vasilarou, Nikolaos Alachiotis, Joanna Garefalaki, Apostolos Beloukas, Pavlos Pavlidis

**Affiliations:** Institute of Molecular Biology and Biotechnology (IMBB), Foundation for Research and Technology Hellas (FORTH); Department of Biology, University of Crete, Crete, Greece; Institute of Computer Science (ICS), Foundation for Research and Technology Hellas (FORTH); Technical University of Crete, Crete, Greece; Department of Biomedical Sciences, University of West Attica, Athens, Greece; Institute of Infection and Global Health, University of Liverpool, Liverpool, UK

**Keywords:** SARS-CoV-2, Population genetics, Recombination, Mutation rate, Selective sweeps, Demographic inference

## Abstract

The current coronavirus disease 2019 (COVID-19) pandemic is caused by the SARS-CoV-2 virus and is still spreading rapidly worldwide. Full-genome-sequence computational analysis of the SARS-CoV-2 genome will allow us to understand the recent evolutionary events and adaptability mechanisms more accurately, as there is still neither effective therapeutic nor prophylactic strategy. In this study, we used population genetics analysis to infer the mutation rate and plausible recombination events that may have contributed to the evolution of the SARS-CoV-2 virus. Furthermore, we localized targets of recent and strong positive selection. The genomic regions that appear to be under positive selection are largely co-localized with regions in which recombination from non-human hosts appeared to have taken place in the past. Our results suggest that the pangolin coronavirus genome may have contributed to the SARS-CoV-2 genome by recombination with the bat coronavirus genome. However, we find evidence for additional recombination events that involve coronavirus genomes from other hosts, i.e., Hedgehog and Sparrow. Even though recombination events within human hosts cannot be directly assessed, due to the high similarity of SARS-CoV-2 genomes, we infer that recombinations may have recently occurred within human hosts using a linkage disequilibrium analysis. In addition, we employed an Approximate Bayesian Computation approach to estimate the parameters of a demographic scenario involving an exponential growth of the size of the SARS-CoV-2 populations that have infected European, Asian and Northern American cohorts, and we demonstrated that a rapid exponential growth in population size can support the observed polymorphism patterns in SARS-CoV-2 genomes.

## Introduction

In late December 2019, Chinese health authorities reported a cluster of atypical pneumonia cases epidemiologically linked with the Huanan Seafood Wholesale Market in Wuhan, Hubei Province, China (Huang et al., 2020). On January 7, 2020, these cases were associated with a novel human coronavirus (hCoV), dubbed SARS-CoV-2 (Coronaviridae Study Group of the International Committee on Taxonomy of Viruses, 2020), which consists the third documented spillover from mammals, but it is divergent from SARS-CoV and MERS-CoV that caused past epidemics (Rota et al., 2003; Zaki et al., 2012). Two weeks later, the United States of America reported the first confirmed case of SARS-CoV-2 infection while the first three cases in Europe were confirmed on January 24, 2020 (Holshue et al., 2020). As of April 20, the pandemic coronavirus-associated acute respiratory disease called coronavirus disease 19 (COVID-19) has infected more than 2,397,216 people and has caused 162,956 deaths (WHO, COVID-19 Situation Report 92).

As the outbreak progresses over time, laboratories around the world are sequencing SARS-CoV-2 genomes from various human sources and timepoints to track the dispersal pattern of the pandemic. SARS-CoV-2 belongs to the Betacoronavirus genera and structurally is an enveloped RNA virus, with a non-segmented, positive-sense (+ssRNA) genome of ~30 kb, amongst the largest identified RNA genomes (Andersen et al., 2020; Zhu et al., 2020). The application of next-generation sequencing (NGS) approaches on pathogens can elucidate important features like disease transmission and virulence (Gwinn et al., 2019). In fact, NGS data are accumulating with unprecedented speed, and the Global Initiative on Sharing All Influenza Data (GISAID) database, which originally promoted the international sharing of all influenza virus sequences, as of April 2, included 3,230 SARS-CoV-2 genome data submissions (2,305 full-length sequences with high-coverage).

Population genetics can provide insights into the spread and epidemics of viruses because it offers the machinery to estimate the values of parameters such as the mutation and recombination rate. Both of these parameters are important for the evolution and the management of viral diseases. Mutation rate, recombination, as well as stochastic (random genetic drift) and non-stochastic processes (selection) are the dominant forces that shape viral diversity in natural populations of RNA viruses. In fact, due to the lack of proofreading in RNA-dependent RNA polymerases, RNA viruses demonstrate the highest mutation rates of any group of organisms (Moya et al., 2004), which may lead to viral adaptation to selective pressures and alter their virulence. In addition, recombination has an important role in evolution, by elevating haplotypic variability in viral populations and potentially generating highly fitted genomes more rapidly than by mutation rate alone, especially if epistatic interactions are present between different genomic locations. Research on viral recombination rates has demonstrated its association with increased genetic diversity and the generation of novel lineages and new pathogenic recombinant circulating viruses (Combelas et al., 2011; Pérez-Losada et al., 2015). Actually, recent studies suggest that recombination events may have shaped the architecture of SARS-CoV-2 genome by demonstrating that it shares 96.3% genetic similarity with a bat CoV (Lu et al., 2020) and 91.2% genetically similarity with two pangolin CoV genomes (Andersen et al., 2020; Boni et al., 2020; Lam et al., 2020; Liu et al., 2019; Paraskevis et al., 2020). High mutation and/or recombination rates imply that developing a successful vaccine will be challenging since viral genomes will be able to adapt fast in human interventions (Andersen et al., 2020). Moreover, population genetics provides tools to infer the recent evolutionary history of populations, both adaptive and non-adaptive. Thus, by using statistical approaches, such as the Approximate Bayesian Computation (Beaumont et al., 2002), it is feasible to assess past population size changes by using only present-day genomic data. RNA viruses demonstrate immense population sizes (Moya et al., 2004) and the study of the population demographic history could support viral epidemiological research. A previous study, in particular, has associated the change from a bat CoV population of constant size to a population growth of CoVs from other hosts, with the interspecies transmission of viruses from their original reservoir to an alternate host (Vijaykrishna et al., 2007). Furthermore, selective sweep theory (Kim & Stephan, 2002; Maynard Smith & Haigh, 1974; Stephan et al., 1992) allows to localize the targets of recent and strong selection on genomes.

In our study, we aim to elucidate the recent evolution of SARS-CoV-2 by implementing population genetics approaches on full-length SARS-CoV-2 genomes available at the GISAID database (https://www.gisaid.org/). Understanding the SARS-CoV-2 evolutionary processes would enable us to assess critical parameters, such as the mutation rate, and plausible recombinations, as well as recent selective sweeps and population demographic parameters, which can directly affect the evolution of this virus within the human population. Such results will support epidemiological research on tracing dispersal patterns of the pandemic and designing therapeutic or preventive strategies.

## Results

We used publicly available data of SARS-CoV-2 genomes from the GISAID database, downloaded on 2/4/2020. All analyses, but selective-sweep localization, were performed on this dataset. The initial dataset comprises 3,230 full genome assembled sequences. From those, we kept only the high-coverage sequences (as defined by GISAID, these sequences contain less than 1% ambiguous states (N’s) and less than 0.05% unique amino acid mutations, i.e., not seen in other sequences in the database, and no insertion-deletions unless verified by submitters). In total, 2,305 pass this filter. We applied additional filters to keep only high-quality sequences: First, we trimmed the ambiguous N’s from both the beginning and the end of the genomes. Then, after trimming, we kept only sequences with less than 10 ambiguous states (N’s). The final dataset consists of 1,895 SARS-CoV-2 genomes and two outgroup sequences: the bat CoV (hCoV-19/bat/Yunnan/RaTG13/2013; Accession ID EPI_ISL_402131) and pangolin CoV genomes (hCoV-19/pangolin/Guangdong/1/2019; Accession ID: EPI_ISL_410721). The dataset was aligned using the MAFFT software v.7.205 (Katoh & Standley, 2013).

### Estimation of mutation rate and divergence from bat CoV

We modeled the divergence from the bat CoV genome as a function of the sequence sampling date. Divergence was estimated using the Kimura80 approach (Kimura, 1980) as implemented in libsequence C++ library (Thornton, 2003). Since the sampling date is known (provided by the GISAID database), we were able to perform a regression analysis of the divergence per site as a function of the sampling date. The slope of the line represents the increase of divergence per site and per day, thus it may be used as a proxy for the mutation rate per site and per day. The estimated mutation rate is 1.87 x 10^−6^ per nucleotide substitution per site per day (Figure 1), which is expected and in concordance with what has been previously reported for SARS-CoV-1 (0.80 to 2.38×10^−3^ per site and per year, i.e., 2.24-6.68 x 10^−6^ per site and per day) and ~10^−6^ per site per cycle for other CoVs (Bar-On et al., 2020; Shen et al., 2020; Zhao et al., 2004).

**Figure 1:**
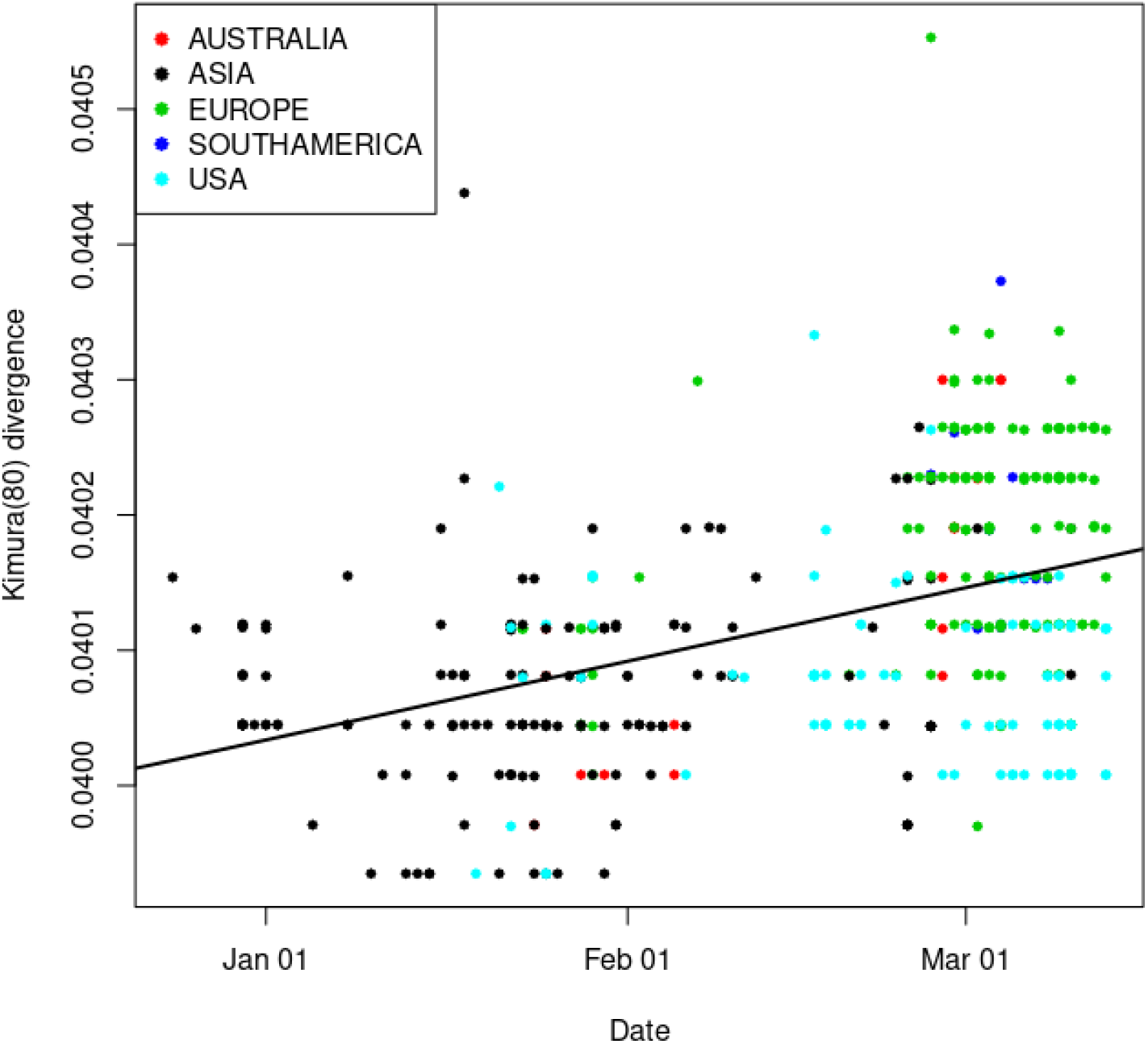
Regression analysis of the divergence from the bat CoV genome as a function of sampling date. The slope of the regression line (1.87 x 10^−6^) reflects the increase in divergence per site and per day.

The divergence from the bat CoV genome is estimated to 0.04 substitutions per site using the Kimura (1980) calculation. Assuming that the mutation rate is 1.87 x 10^−6^, the total time since that separates SARS-CoV-2 from the bat CoV genome is 0.04/1.87 x 10^−6^ = 58.6 years. Thus, assuming an equal mutation rate between the bat CoV lineage and the human CoV genome, the time since the common ancestor between the bat CoV and the SARS-CoV-2 is about 58.6/2 = 29.3 years.

### Recombination events

#### Host analysis

We performed a BLAST-based analysis using as a (locally constructed) database the 55 fully sequenced Coronaviridae genomes downloaded from NCBI (https://www.ncbi.nlm.nih.gov/genomes/GenomesGroup.cgi?taxid=11118) together with the bat CoV and pangolin CoV genomes from the GISAID database. The query comprises all the unique 300-mers from the 1,895 SARS-CoV-2 genomes. In total, there are 452,212 unique 300-mers. The word size parameter of BLAST was set to 10 and the e-value to 0.1 to capture even small parts of the sequence that may resemble parts from the Coronaviridae family of different hosts. We checked whether consecutive regions in a query sequence could match different entries in the local blast database, that is Coronaviridae sequences from different hosts. Then, we estimated all correlation coefficients for all host pairs to evaluate their tendency to co-occur as a match in the same query sequence (Figure 2). For example, pangolin and bat co-occur in 395 out of 5568 total cases, resulting in an *r* value of 0.4. Such a high co-occurrence may suggest that an ancestral sequence of a query 300-mer could have experienced a recombination event in these two hosts. We found plausible past recombination events between CoVs from pangolin and bat, Sparrow and *Rhinolophus blasii* (bat), and the hCoV-HKU1 genome with Hedgehog (*Erinaceus*).

**Figure 2:**
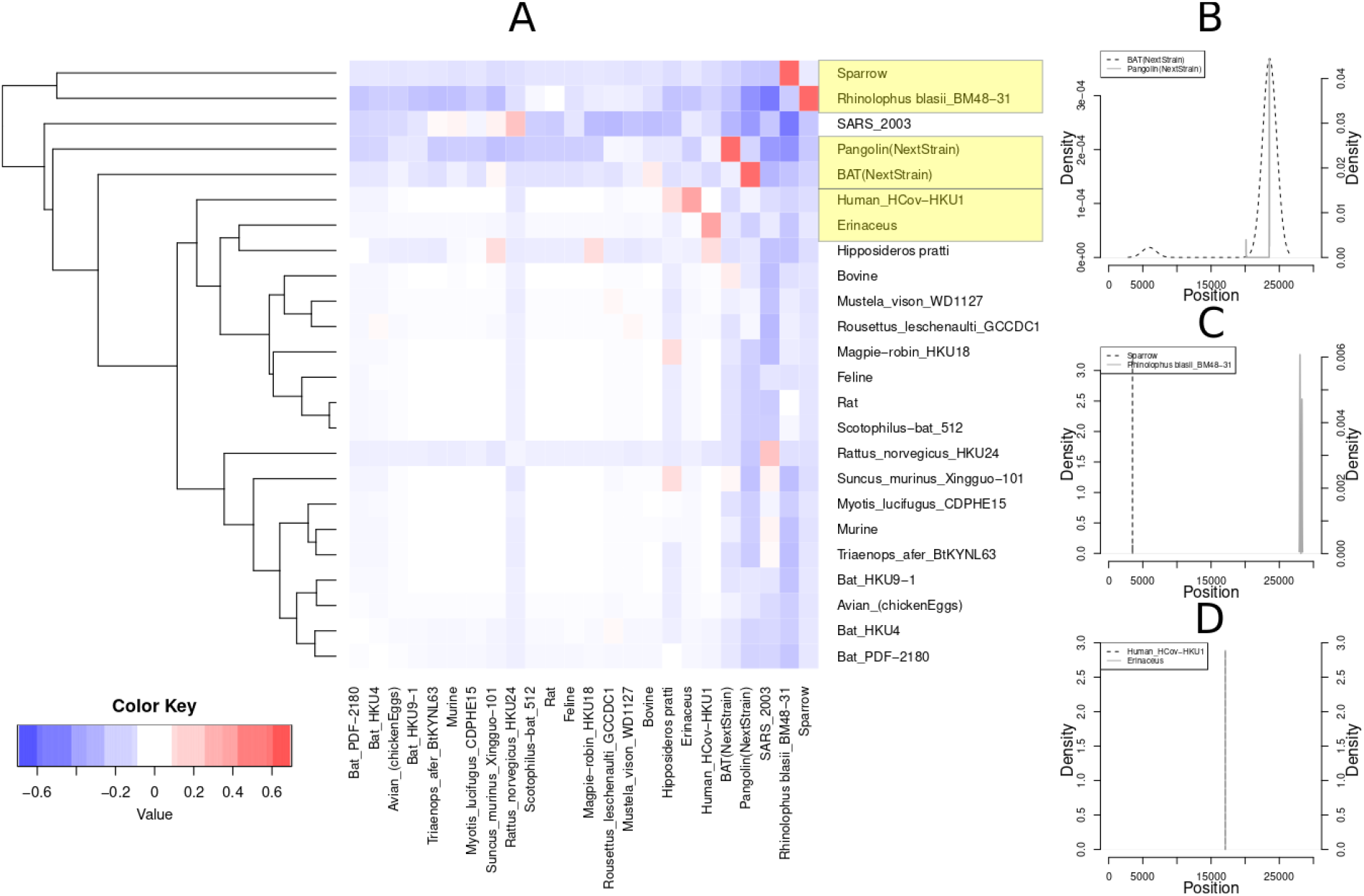
(A) Hosts of Coronaviridae virus family in which plausible recombination events may have taken place. Red-colored squares denote positive correlation coefficient values, thus co-occurence of BLAST results from different hosts for the same query sequence, suggesting potential recombination events. Blue and white squares denote pairs with negative or close to zero correlation coefficient. Yellow boxes denote the host pairs with the highest *r* values: pangolin (GISAID Accession ID EPI_ISL_402131) and bat (GISAID Accession ID: EPI_ISL_410721), Sparrow (RefSeq ID: NC_016992.1) and *Rhinolophus blasii* (RefSeq ID: NC_014470.1), and the Human Cov sequence (RefSeq ID: NC_006577.2) with Hedgehog (RefSeq ID: NC_006577.2). Localization of recombination events in the (B) bat/pangolin pair, (C) Sparrow/Rhinolophus pair and (D) Hedgehog/Human_CoV-HKU1.

**Figure 3:**
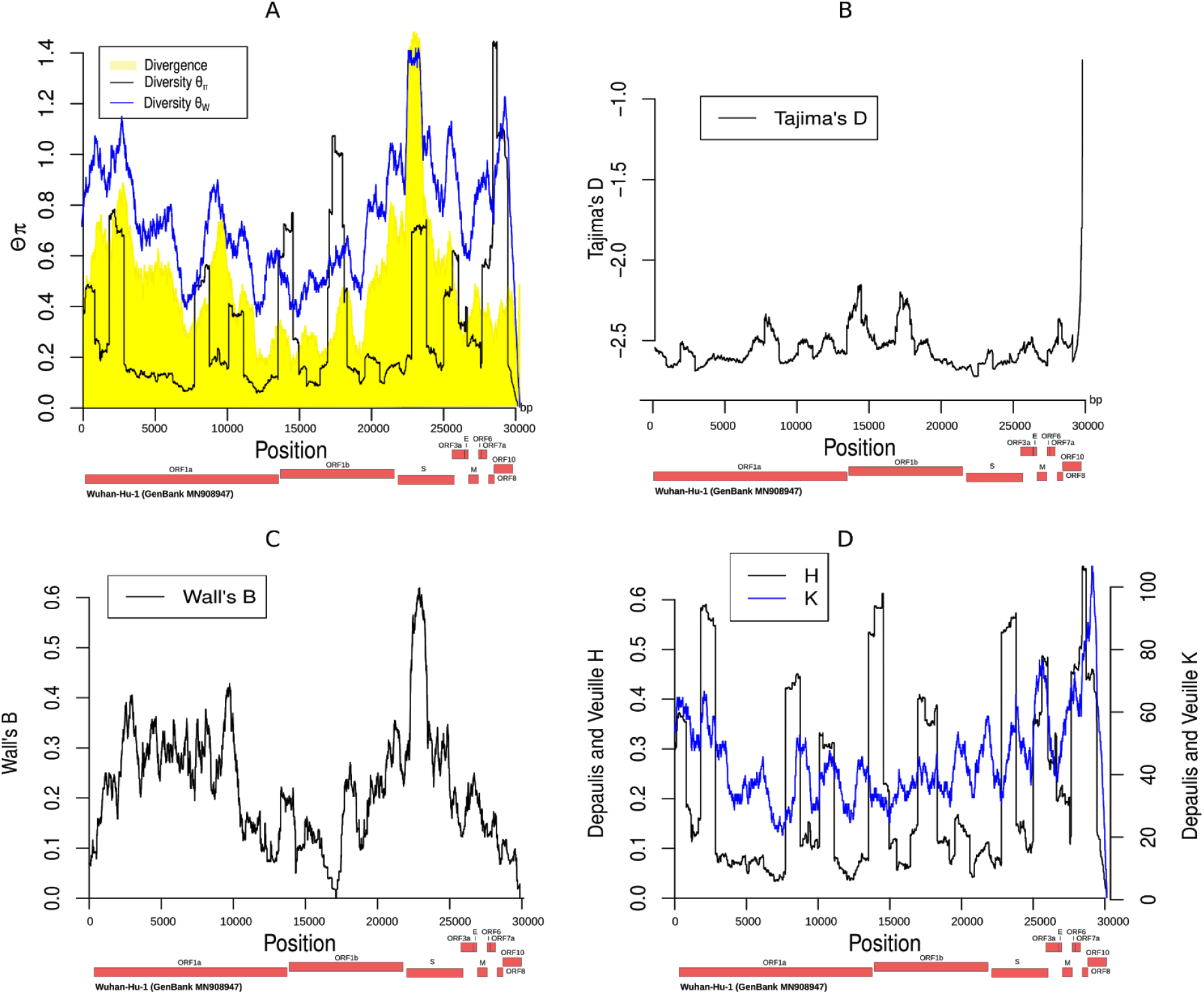
Distribution along the genome of the summary statistics θ_π_, θ_W_, divergence [A], Tajima’s D [B], Wall’s B [C], and Depaulis and Veuille H and K [D]. Wall’s Q is similar to Wall’s B and is not shown in the figure.

#### Localization of recombination events

For the three pairs of hosts depicted in Figure 2, we localized the genomic regions that may be involved in recombination events. Thus, for all concordant pairs, that is pairs corresponding to consecutive matches between a query 300-mer and two different database entries, we used the genomic coordinates of the non-SARS-CoV-2 host. The density plots in Figures 2B, C and D illustrate the distribution of the location in the genome of the host. For the pangolin/bat pair (Figure 2B), the location of both pangolin and bat fragments are around the position 23,583, suggesting a plausible recombination event in the highly divergent Spike protein S1 coding region. On the other hand, the Sparrow/Rhinolophus pair shows different localization of the Sparrow and Rhinolophus fragments. The Sparrow fragments are located around the location 3,484, whereas the Rhinolophus fragments around the position 28,034. Thus, it is probable that an ectopic recombination event between ORF1a and ORF8 has taken place, since it is common for viruses to exchange genetic material in a non-reciprocal manner (Pérez-Losada et al., 2015) even though we cannot exclude the possibility of a false positive result. Finally, the Hedgehog/Human_HCoV-HKU1 pair both locations are around the position 17,000 (17,062 and 17,088, respectively), within the ORF1b polyprotein and the helicase/nsp13 coding region.

#### Detection of recombination events amongst SARS-CoV-2 genomes

Using a similar approach as in “Host analysis”, we assessed whether recombination events have taken place between the human SARS-CoV-2 genomes. No recombination event was detected using the SARS-CoV-2 sequences from the GISAID database until the date 2/4/2020.

### Linkage Disequilibrium (LD) analysis

An indirect approach to infer recombination in a sequence is the decay of LD as a function of distance. We estimated LD between all sites using the software *plink* (Purcell et al., 2007) and we modeled haplotypic LD as a function of distance between two genomic sites using a simple linear model as LD = *a*x + *b*, where x is the distance. The analysis was performed on the SARS-CoV-2 genomes for each population separately (Europe, Africa, North America, South America, Oceania, Asia). Results for parameters *a* and *b* are presented in the Supplementary Table S1. These results suggest that for the European, Asian and the total population, LD is decreasing as a function of distance. Thus, recombination may have occurred in SARS-CoV-2 genomes. These findings are not contradictory to the previous section “Detection of recombination events between the SARS-CoV-2 genomes”. In the previous section, our goal was to detect the recombination breakpoints and the recombinant sequences (or the descendants of the recombinant sequences). However, if the sequences are not divergent enough such an approach will not reveal the breakpoints. On the contrary, the LD decay approach is more sensitive to show that recombination may have happened (however, it can neither reveal the recombination breakpoints nor the recombinant sequences). The Supplementary Table S1 shows that in the North America, South America, Oceania and Africa populations, the value of the slope *a* is not statistically different than 0, i.e., there is no correlation between the genomic distance and the LD. This is probably due to the fact that the number of recombination events that take place in a population is a function of both the recombination rate and the effective population size (ρ = 4N_e_r, where N_e_ is the effective population size and r the recombination rate). Since, the exponential growth phase started later in these continents compared to Asia and Europe, it is possible that the effective population size of the virus was not large enough for recombination to be detected in the North America, South America, Oceania and Africa populations.

### Selective sweeps

Selective sweep analyses were done on the dataset downloaded on 31/3/2020 due to extensive computational time requirements. We used SweeD (Pavlidis et al., 2013) and RAiSD (Alachiotis & Pavlidis, 2018) to identify potential targets of selective sweeps. SweeD implements a composite likelihood ratio test based on the site-frequency spectrum, while RAiSD evaluates the μ statistic that relies on multiple sweep signatures. Supplementary Figure S1 illustrates the scores of both tools along the genome when analyzing 1,601 SARS-CoV-2 sequences (version 31/3/2020) in the total population and per continent for Europe, Asia, and North America. The figure also shows the top 5% common outliers. Given the accuracy granularity of existing sweep detection methods, including the employed ones in this study, we refrain from identifying specific genomic positions at site granularity.

Table 1 provides distinct region pairs that were identified as potential sweep targets. For the world sample, the first two region pairs are located within the non-structural protein 3 (nsp3), which is a large multidomain and multifunctional protein that plays a key role in coronavirus replication. Previous studies report that the nsp3 represents a preferential selection target in MERS-CoV and lineage C betaCoVs (Forni et al., 2016). The third region pair belongs to Spike protein S1, which attaches the virion to the cell membrane by interacting with a host receptor, thereby initiating the infection. The highly divergent Spike protein S1 of coronaviruses facilitates viral attachment, fusion and entry, and thus serves as a target for antibody and vaccine development (Tai et al., 2020). The identified outlier region 23,043-23,073 falls into the receptor-binding domain (RBD) which has been reported to be optimized, as a result of natural selection, for binding to human angiotensin-converting enzyme 2 (ACE2) receptors and consists a putative target for designing future therapeutic or preventive strategies (Andersen et al., 2020). The fourth region pair is in the spike protein S2, which mediates the fusion of the virion and cellular membranes by acting as a class I viral-fusion protein.

**Table 1:**
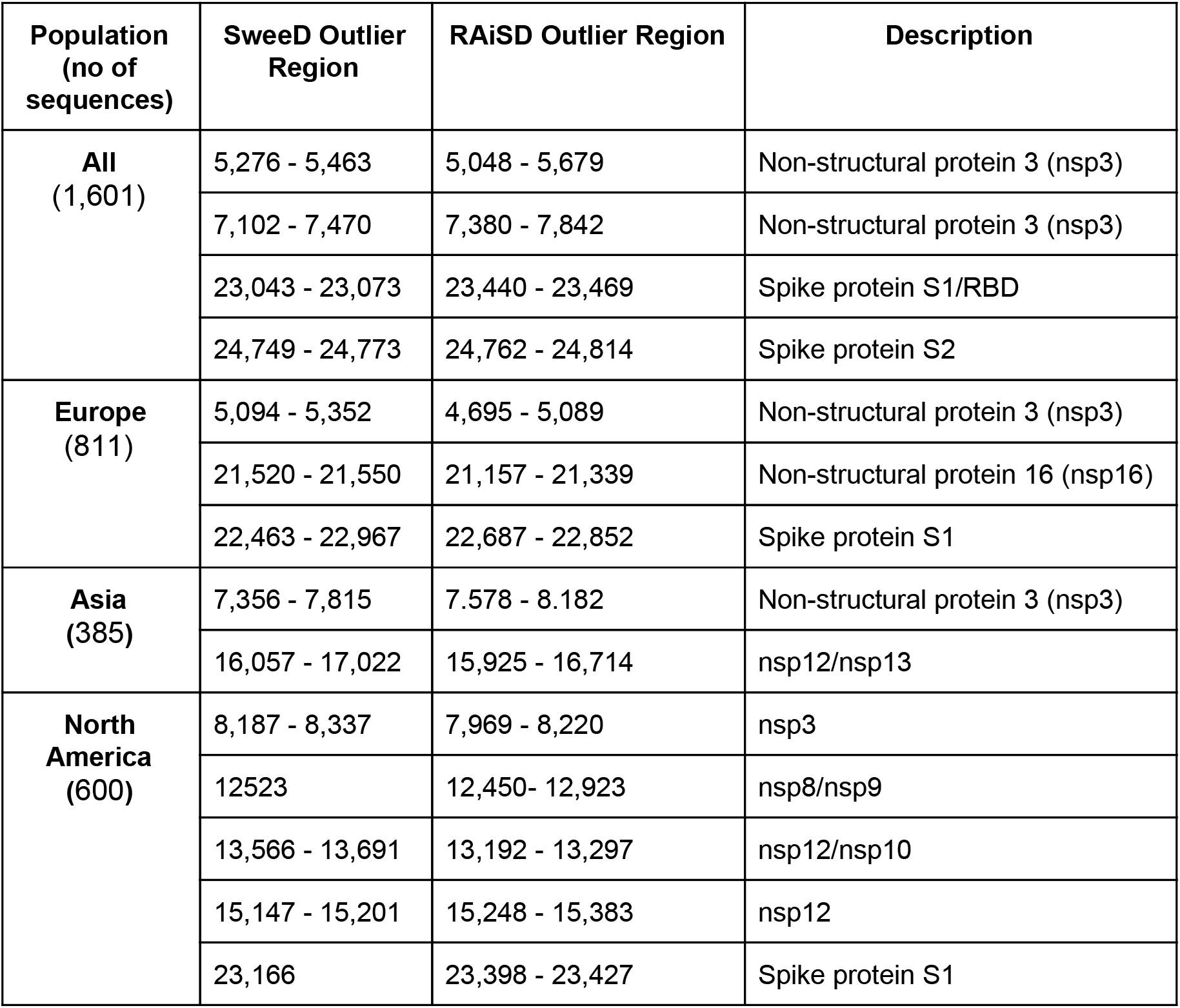
Common-outlier region pairs between SweeD and RAiSD based on a maximal distance of 0.4kb.

| Population<br>(no of<br>sequences) | SweeD Outlier<br>Region | RAI SD Outlier Region | Description |
| --- | --- | --- | --- |
| **All**<br>(1,601) | 5,276 - 5,463 | 5,048 - 5,679 | Non-structural protein 3 (nsp3) |
|  | 7,102 - 7,470 | 7,380 - 7,842 | Non-structural protein 3 (nsp3) |
|  | 23,043 - 23,073 | 23,440 - 23,469 | Spike protein S1/RBD |
|  | 24,749 - 24,773 | 24,762 - 24,814 | Spike protein S2 |
| **Europe**<br>(811) | 5,094 - 5,352 | 4,695 - 5,089 | Non-structural protein 3 (nsp3) |
|  | 21,520 - 21,550 | 21,157 - 21,339 | Non-structural protein 16 (nsp16) |
|  | 22,463 - 22,967 | 22,687 - 22,852 | Spike protein S1 |
| **Asia**<br>(385) | 7,356 - 7,815 | 7.578 - 8.182 | Non-structural protein 3 (nsp3) |
|  | 16,057 - 17,022 | 15,925 - 16,714 | nsp12/nsp13 |
| **North America**<br>(600) | 8,187 - 8,337 | 7,969 - 8,220 | nsp3 |
|  | 12523 | 12,450- 12,923 | nsp8/nsp9 |
|  | 13,566 - 13,691 | 13,192 - 13,297 | nsp12/nsp10 |
|  | 15,147 - 15,201 | 15,248 - 15,383 | nsp12 |
|  | 23,166 | 23,398 - 23,427 | Spike protein S1 |

### Summary statistics along the SARS-CoV-2 genome

We estimated the values of several summary statistics, commonly used in population genetics studies, along the genome of SARS-CoV-2: (i) Watterson’s estimate (Watterson, 1975) of θ value (the population mutation rate); (ii) θ_π_ (Tajima, 1983), i.e., the average number of pairwise differences in the sampled sequences, which is another estimate of θ; (iii) Tajima’s D (Tajima, 1989), a commonly used neutrality test that receives negative values typically in cases of population expansion and/or recent strong positive selection; (iv) Wall’s B and Wall’s Q (Wall, 1999), which are related to the recombination rate; (v) the divergence to the bat sequence using a random SARS-CoV-2 genome, the average divergence from the bat Coronavirus sequence using all SARS-CoV-2 genomes and (vi) the Depaulis and Veuille K and H statistics (Depaulis & Veuille, 1998), which denote the number of haplotypes and the haplotype heterozygosity, respectively. The region around position 23,000 (Spike protein S1/RBD) is characterized by large divergence, low Tajima’s D, and high values of Wall’s B and Q. The divergence values (high) and the Tajima’s D values (low) suggest that it may be a target of recent positive selection. The value of θ_W_ is high, which is due to the excessive number of singletons in the region since θ_π_ does not receive very high values. This result is consistent with positive selection, even though we cannot exclude artifacts from sequencing errors or more complex evolutionary phenomena (e.g., recombination with another host). Another probable explanation is that selective sweep is ongoing. Thus, there will still exist several low-frequency polymorphisms decreasing Tajima’s D and maintaining high values for θ_W_.

### Estimation of the time of the most recent common ancestor

We used all samples to estimate the mean and the variance of the parameter *t* (Supplementary Figure S2). The mean parameter value estimation was *t_est_* = 47.0 and the variance 504.45. Supplementary Figure S2B, depicts the histogram, density and the mean value for the parameter *t*, the time of the most recent common ancestor measured in days before the first Wuhan sample (24/12/2019). The estimated value of *t* suggests that the most recent common ancestor of SARS-CoV-2 genomes is quite older (47 days) than the first reported case. The estimated value is similar, though a bit older, to the value presented in NextStrain (www.nextstrain.org) using a regression analysis (NextStrain dates the TMRCA on 14/12/2019 as estimated on April 21). Even though it is reasonable that the first case is dated earlier than the first reported case, here, we cannot know whether the TMRCA refers to a human sample or another host (e.g., bat).

### Demographic inference

Exploring the demographic dynamics of SARS-CoV-2 populations is important in understanding the magnitude of the spread of the ongoing COVID-19 pandemic. Here, we focused on the SARS-CoV-2 sequences of Asian, European and Northern American human cohorts and we employed an Approximate Bayesian Computation (ABC) approach to model the exponential growth of the three SARS-CoV-2 populations. For each population, the parameter inference procedure involves the estimation of the posterior distributions of two parameters: the scaled population mutation rate *θ* and the rate of exponential expansion *α*. In this model of population growth the population size is given by: N(t) = N_0_ e^−αt^, where time t is measured backwards in time in units of 2N_0_ generations and *θ* is defined as 2N_0_μ for haploid organisms, where μ is the mutation rate per generation and N_0_ the present-day effective population size (Hein et al., 2005).

For each population under study, ABC employs stochastic coalescent simulations to estimate properties of the parameters’ posterior distributions. The parameter inference procedure consists in retaining only the best τ of the simulations, i.e., τ of the simulated datasets which are closest to the observation (Supplementary Figure S3–5). Visualizing in a contour plot the percentage of accepted simulations that was determined with a tolerance τ=0.005 (Supplementary Figure S6–8), demonstrates how these simulations explore the space around the observed data. The summary statistics form an 8-dimensional space. For some statistics (i.e., dimensions), the contour lines diverge from the observed value, because the algorithm keeps the simulations that minimize the 8-dimensional Euclidean distance to the observed vector. Thus, even if the distance is minimized over all dimensions, this may not be true for each dimension separately. For example, the posterior distributions of Tajima’s D and DVK of the European population in particular, show that the accepted values (within the tolerance τ) are quite distant from the observed value. Specifically Tajma’s D and DVK are affected by low frequency derived variants, especially singletons. Tajima’s D becomes negative and DVK assumes large values in the presence of singletons since the number of haplotypes increases. In the SARS-CoV-2 dataset a large proportion of polymorphisms (>70%) are singletons, which results in extreme negative values of Tajima’s D. Such a proportion of singletons is difficult to be explained with a simple demographic model of population expansion alone. It is possible that other factors (e.g., sequencing errors, more complex demographic models) have contributed to such an observation.

An improvement to the simple rejection ABC algorithm was employed to correct for the discrepancy between the simulated and the observed statistics. The posterior probability of the parameters was approximated by applying regression adjustment techniques to the retained simulations. More specifically, we performed a local linear weighted regression correction to estimate the posterior parameter probabilities of the American and the Asian SARS-CoV-2 population while a non-linear regression model using feed-forward neural networks was utilized in the demographic analysis of the European population. This correction influenced our estimates, reducing the variance of the posterior estimates (Figure 4). The inferred mean, median and mode values, as well as quantile values are shown in Table 2. Both α and θ values are higher in the Asian and North American populations than the European population. This suggests that population growth was faster in Asia and North America compared to the European population. Also, since θ = 2N_0_μ (N_0_ is the present day population size for each population) is higher for the Asian and North American population, it suggests that N_0_ is higher in Asia and North America than Europe. We note that N_0_ is different from the effective population size: the effective population size N_e_ represents a long-term ideal population size, while N_0_ refers only to the present. Furthermore, we note that *a-posteriori* the two parameters α and θ are correlated (Supplementary Figure S9). Thus, the likelihood surface may have multiple local optima and the inference process may result in different optima (and therefore, parameter estimates) in different populations.

**Figure 4:**
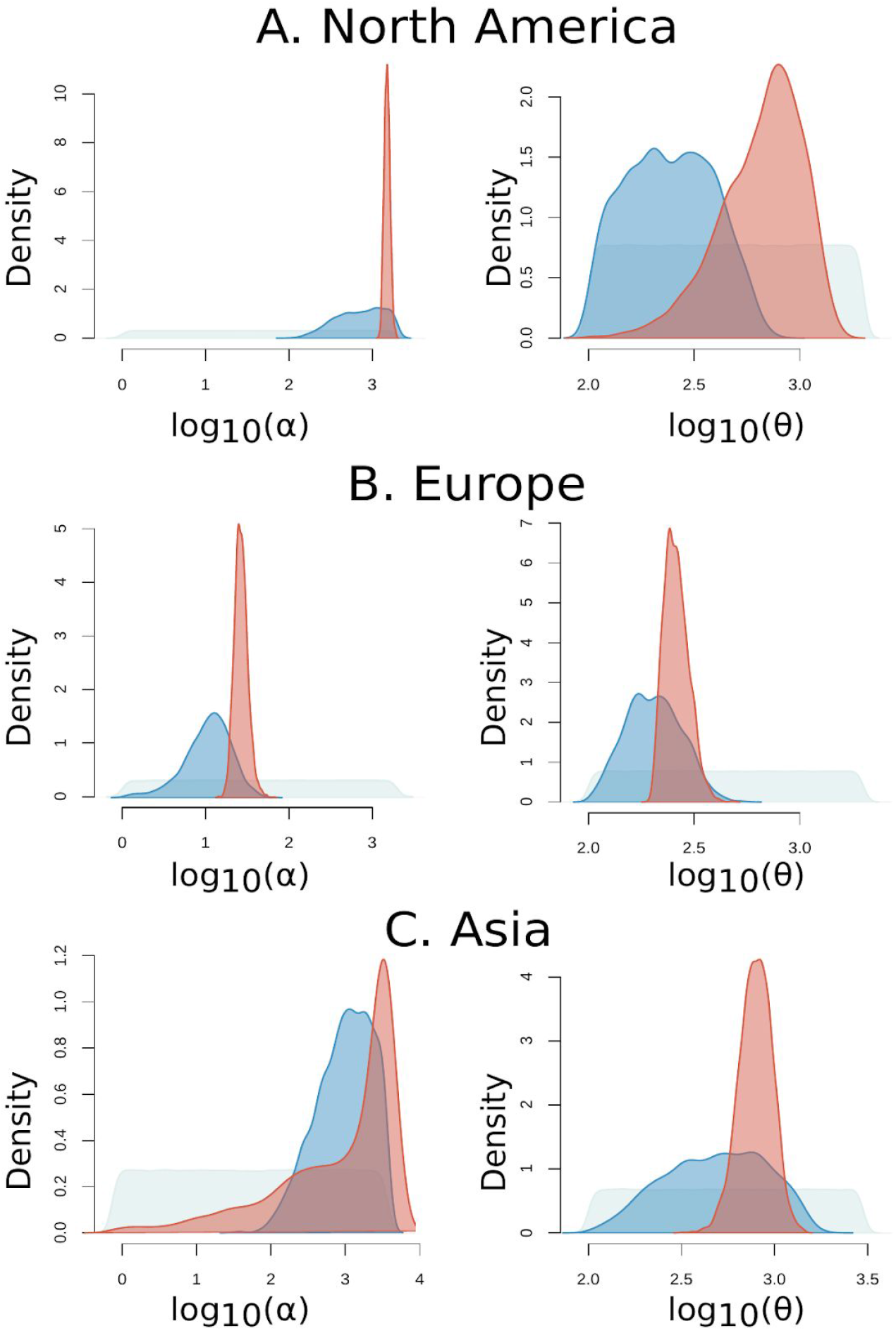
Probability estimations of (A) North America’s, (B) Europe’s and (C) Asia’s SARS-CoV-2 population expansion parameters. Grey curve: Prior parameter distribution; Blue curve: Density distribution of the ABC unadjusted simulation values that were accepted with a tolerance of 0.005; Red curve: Posterior distribution approximated by applying regression correction to the retained simulations.

**Table 2:** Summary statistics of the posterior distributions that were estimated by ABC for each SARS-CoV-2 population.

|  | North America |  | Asia |  | Europe |  |
| --- | --- | --- | --- | --- | --- | --- |
| | $\alpha$ | $\theta$ | $\alpha$ | $\theta$ | $\alpha$ | $\theta$ |
| **Weighted Mean 2.5% Percentile** | 1299.12 | 235.37 | 12.91 | 523.95 | 19.26 | 211.95 |
| **Weighted Median** | 1500.96 | 709.24 | 2310.46 | 794.61 | 26.21 | 255.9 |
| **Weighted Mean** | 1509.05 | 717.89 | 2508.8 | 807.84 | 26.85 | 260.3 |
| **Weighted Mode** | 1507.6 | 733.87 | 4869.54 | 738.66 | 24.81 | 241.78 |
| **Weighted Mean 97.5% Percentile** | 1766.13 | 1300.2 | 4999.94 | 1147.67 | 38.63 | 333.24 |

## Methods

### Mutation rate analysis and estimation of the time of the most recent common ancestor between bat CoV and SARS-CoV-2

Let μ the mutation rate per nucleotide and per day. The collection date of the samples has been recorded in the GISAID repository. Let *h_i_* be the collection date for sample *i*. If we perform a regression analysis of the number of nucleotide differences (divergence) *div_i_* between sample *i* and the bat Coronovirus sequence as a function of collection date, there should be a relation of the form: *div_i_* = *h_i_* * μ + β, i.e., the divergence between sample *i* and the bat Coronavirus increases linearly with time. The parameter β denotes the intercept, i.e., the divergence of the first sampled SARS-CoV-2 sequence in Wuhan. To estimate the parameter μ, we calculated the slope of the regression line that models divergence as a function of sampling date. Furthermore, the point that this line crosses the x-axis (the axis of dates), denotes the time point t_bat/Cov2_ at which the divergence between bat and SARS-CoV-2 is 0, i.e., the time of the most recent common ancestor between bat and SARS-CoV-2.

### Recombination analysis

Recombination evaluation was performed with a blast-based analysis. Let D_S_ (Dataset Sars) denote the SARS-CoV-2 fasta dataset as it was downloaded from the GISAID database. Furthermore, D_C_ (Dataset Coronavirus) the dataset with 55 Coronaviridae sequences downloaded from NCBI and with the pangolin and bat sequences from GISAID (thus, 57 sequences in total), and D_BP_ (Dataset pangolin/bat) is a dataset with only two sequences, the bat and the pangolin sequence from GISAID. First, using D_S_, we generated a new dataset called D_SS_ (Dataset Sars Split) which comprises all the unique 300-mers from the *whole* D_S_. To detect potential contribution of bat/pangolin recombinant sequences into SARS-CoV-2 we blasted the D_SS_ dataset against the D_BP_ dataset (i.e., D_BP_ is the blast database and D_SS_ the query set). If a sequence D_SS,i_ (or an ancestral sequence of it) is the result of recombination between the bat Coronavirus sequence and the pangolin Coronavirus sequence, then the sequence will be matched to both bat and pangolin in consecutive parts of it. Thus, if D_SS,i_ is a sequence of length L (here L = 300bp), we are looking for blast results of the the form: D_SS,i_ [*a*, *b*] matches to bat/pangolin and D_SS,i_[c, d] matches to pangolin/bat, where 1 ≤ *a* ≤ *b* ≤ *c* ≤ *d* ≤ L. We applied also the following restrictions: 1 ≤ *a* ≤ 10, L-10 ≤ *d*≤ L, |*b* - *c*| ≤ 20. In other words, *a* should be in the beginning of the 300-mer, *d* should be at the end of the 300-mer and the distance between the bat-like and the pangolin-like parts should be smaller than 20. To detect other potential recombination events, we used as a blast database the D_C_ dataset to search for recombinants from other hosts besides bat and pangolin. Furthermore, to detect recombination between human SARS-CoV-2 sequences, or their unsampled ancestral sequences, we used as a blast database the D_S_ dataset. The specific parameters for running *blastn* were the following: -outfmt 6 (to present the results in a tabulated format), -word_size 10 (to capture even small fragments) and -evalue 0.1 (also, to be able to capture small fragments).

### Linkage Disequilibrium (LD)

An indirect approach to infer recombination in a sequence is the decay of LD as a function of distance. Given the demographic model, LD between two sites may be decreased either due to recombination events or due to recurrent mutations, violating, thus, the infinite site model (Kimura, 1969). From these two causal reasons of LD decay, only recombination decreases LD as a function of the distance between two sites since the recombination rate is proportional to the distance of the two sites. We used *plink* to estimate haplotypic LD between all possible pairs of polymorphisms in the SARS-CoV-2 genome. Then, we modeled LD as a linear function of distance (assuming that the recombination rate is constant along the genome) as follows:

LD = *ax* + *b*, where *x* is the distance between two sites and *a* and *b* the slope and intercept, respectively.

If *a* is negative, then it supports the presence of recombination on the SARS-CoV-2 genome. On the other hand, if *a* is not statistically different than 0, then recombination may not be a reason for LD patterns along the genome.

### Selective sweeps and common outliers

To estimate potential targets of selective sweeps, we deployed the software tools SweeD (Pavlidis et al., 2013) and RAiSD (Alachiotis & Pavlidis, 2018). A selective sweep analysis detects characteristic patterns of polymorphisms attributed to the action of recent and strong positive selection from a new allele. The key mechanism that generates such polymorphic patterns is the fast increase of the beneficial mutation and the recombination rate (Pavlidis et al., 2008). Currently, the whole population of SARS-CoV-2 is rapidly expanding. Furthermore, even though recombination may be present in viruses, our analyses did not find evidence of recombination among human SARS-CoV-2 sequences. This outcome does not suggest that recombination is absent, but rather that diversity within the human SARS-CoV-2 is not large enough to allow the detection of recombination events. Thus, a selective sweep analysis may still be meaningful for SARS-CoV-2.

We performed a selective sweep analysis for each separate population (Asia, Europe, North America, South America, Oceania, Africa) and for the total worldwide sample. SweeD was executed using its default parameters and a grid size of 5,000 positions along the genome (-grid parameter), which led to the calculation of a CLR score every 6 nucleotides, on average. Due to the tool’s high computational complexity for computing the site frequency spectrum, which can lead to prohibitively long execution times when the sample size increases, we used a smaller sample size for the worldwide analysis. The SweeD-based analysis of the employed dataset, which comprised 1,601 genomes, required 112 CPU hours. Initial analyses with smaller sample sizes (performed as data were becoming available online for a period of three weeks) revealed that SweeD results remain largely identical when the sample size exceeds 1,000 sequences.

Unlike SweeD, which relies on maximum-likelihood estimation at predefined positions along the data, RAiSD computes the μ statistic using a polymorphism-driven sliding-window algorithm. RAiSD was executed using a sliding-window size of 8 polymorphisms (-w parameter) and a step of 1 polymorphism (default). We also increased the slack for the SFS edges to 2 (-c parameter), which directed the tool to also consider doubletons and polymorphisms that belong to the S-2 class in the calculation of the μSFS factor, where S is the sample size. The default value for the SFS edge slack is 1, considering only singletons and polymorphisms in the S-1 class.

To conduct a common-outlier analysis based on the results obtained by SweeD and RAiSD, we implemented a series of extensions directly in the source code of RAiSD. More specifically, we introduced a new parameter (-G) that specifies the grid size similarly to SweeD, which directs RAiSD to report scores at the same positions as SweeD when the same grid size is given. Given the RAiSD polymorphism-driven approach, however, scores for the required positions are calculated using linear interpolation based on the initially evaluated positions. To detect common-outlier regions, we sort the evaluated positions in each of the tool reports based on their associated scores, and set the top 0.05 value as the cutoff threshold per method. This allowed us to identify all outlier positions per tool. Thereafter, we compute all pairwise base distances between SweeD and RAiSD outlier positions, and report candidate outlier position pairs if the evaluated positions are closer than 400 bases. The common-outlier analysis was integrated as a feature in RAiSD to facilitate reproduction of the results. The following example command line was used to detect common-outlier regions:

RAiSD -I inputFile -n runName -CO SweeDReport 1 2 -COT 0.05 -O -COD 400 -c 2 -w 8 -G 5000

The -CO parameter provides a path to the SweeD report, as well as the column indices for the positions and the scores. The -COT and -COD parameters specify the cutoff threshold for the outliers and the maximum distance between SweeD/RAiSD outlier positions, respectively.

### Estimation of the time of the most recent common ancestor

To estimate the time of the most recent common ancestor, we applied the following: Let x_0_ be the sequence of SARS-CoV-2 from the first reported patient in Wuhan. This sampling/collection date is 24/12/2019. Any other sequence y_i_ will have a common ancestor with x_0_ at some time t_i_ as shown in Supplementary Figure 1, earlier than the date 24/12/2019. To assess t_i_ we need to consider the differences (diversity), δ_i_, between x_0_ and y_i_. The older t_i_ is, the more differences on average will exist between x_0_ and y_i_. In general, the expected number of differences, δ_i_, between x_0_ and y_i_ will be δ_i_ = (D_i_ + 2t_i_)μ*l*, where μ is the mutation rate per base per day, *l* is the length of the genome (here we used the value 30,000bp), and D_i_ the number of days between the sampling date of x_0_ and y_i_ Substituting δ_i_ with the observed number of pairwise differences between x_0_ and y_i_, μ = 1.87 x 10^−6^ (see “Estimation of mutation rate and divergence from bat” in Results), we can estimate t_i_ for sample *i*. Finally, we estimated the expected [t_i_] and the variance V_t_ of t_i_ as the average of t_i_’s, and the sample variance, respectively. The reason behind using the first reported patient in Wuhan is that the common ancestor between this sequence and any other sequence should be either on 24/12/2019 or earlier than this date. Contrary, it is possible that any two different sequences y_i_ and y_j_ (sampled later during the pandemia) have a common ancestor more recently than the date 24/12/2020, thus it would not be possible to use them in order to estimate the time of the most recent common ancestor.

### Demographic inference

Demographic analysis was performed in an approximate Bayesian computation (ABC) framework. ABC is a bayesian approach that bypasses the exact likelihood computation by using stochastic simulations and summary statistics. (Beaumont et al., 2002). In our analysis, we are modeling the demography of the SARS-CoV-2 dataset that has infected Asian, European and Northern American cohorts. An important approximation in an ABC analysis, is the replacement of the full data by a set of summary statistics. These are numerical values calculated from the data, so that they represent the maximum amount of information in the simplest possible form (Csilléry et al., 2010). The summary statistics were calculated from the multiple sequence alignment (MSA) of each population’s sequences by msABC (Pavlidis et al., 2010) and include (1) estimates of genetic diversity: the Watterson’s estimator *θw* (Watterson, 1975) and the mean pairwise differences of sequences *θπ* (Tajima, 1983), (2) summary of the site frequency spectrum in the form of Tajima’s D (1989), (3) the average pairwise correlation coefficient ZnS (Kelly, 1997) as a measure of linkage disequilibrium, (4) two haplotype-based statistics (Depaulis & Veuille, 1998): the number of haplotypes calculated by the Depaulis and Veuille K (*DVK*) and the haplotype diversity measured by the Depaulis and Veuille H (*DVH*) and (5) the Thomson estimator of the time of the most recent common ancestor TMRCA (Thomson et al., 2000) and its variance (Hudson, 2007).

In order to estimate the parameters of a demographic scenario involving an exponential growth of the population size, we created simulations of neutral polymorphism data. Inferring the demographic history of one population at a time, we generated sets of 500,000 coalescent simulations using msABC (Pavlidis et al., 2010). In that framework, every evolutionary scenario was defined by a set of parameters and every parameter was characterized by a prior distribution (Table 3). Given the current effective size of a population N_0_, the expansion model is characterized by two parameters: the scaled population mutation rate θ and the rate of exponential expansion α. Therefore, the population size is given by: N(t) = N_0_ e^−αt^, where time t is measured backwards in time in units of 4N_0_ generations and θ is defined as 2N_0_μ for haploid organisms, where μ is the mutation rate per generation (Hein et al., 2005). Since there was no clear evidence that recombination has taken place within SARS-CoV-2 genomes, the population recombination rate ρ of all the simulations was set to zero. For each population, the number of individuals sampled per simulation was equal to the number of sequences in the multiple sequence alignment.

**Table 3.**
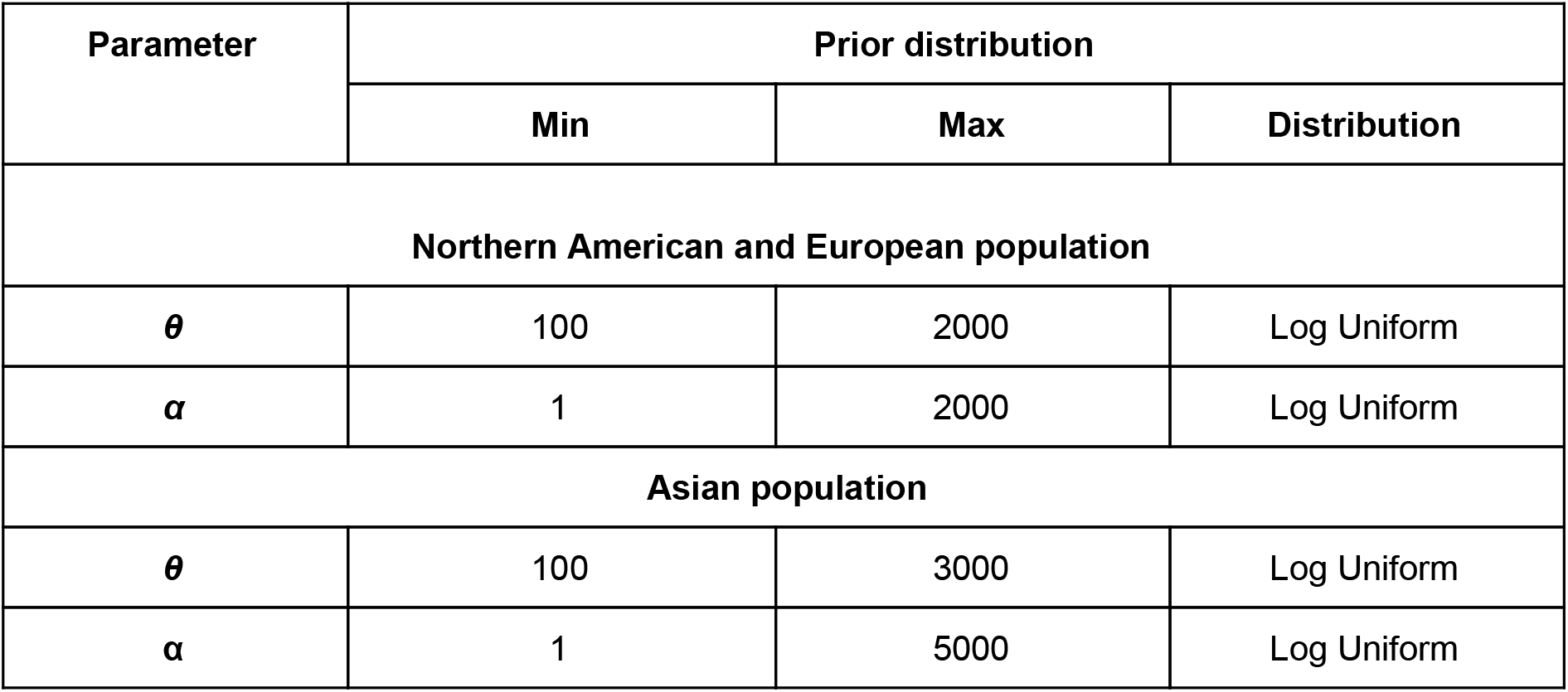
Prior distributions for the demographic parameters.

| Parameter | Prior distribution |  |  |
| --- | --- | --- | --- |
|  | Min | Max | Distribution |
| **Northern American and European population** |  |  |  |
| $\theta$ | 100 | 2000 | Log Uniform |
| $\alpha$ | 1 | 2000 | Log Uniform |
| **Asian population** |  |  |  |
| $\theta$ | 100 | 3000 | Log Uniform |
| $\alpha$ | 1 | 5000 | Log Uniform |

The ABC parameter inference was implemented using the R package abc (Csilléry et al., 2012). The inference procedure consists in retaining simulations for which the Euclidean distance between the set of simulated summary statistics and the observed set is sufficiently small. The percentage of accepted simulations is determined by the tolerance value, τ=0.005. In order to correct for the discrepancy between the simulated and the observed statistics, due to the non-zero τ, the posterior probability of the parameters was approximated by applying regression adjustment techniques to the retained simulations. More specifically, we performed a local linear weighted regression correction to estimate the posterior parameter probabilities of the American and Asian SARS-CoV-2 populations. In this approach, the simulated parameters are assigned weights according to how well the corresponding summary statistics adhere to the observed ones and then a linear regression adjustment is performed to the accepted, weighted parameters (Beaumont et al., 2002). The parameters were ‘logit’ transformed prior to estimation, using as boundaries the prior distributions’ minimum and maximum values that were used during simulations for each parameter. After the regression estimation, the parameters were back-transformed to their original scale. A non-linear heteroscedastic regression model using a feed-forward neural network model was utilized in the demographic analysis of the European population (Blum & François, 2010).

## Acknowledgments

We gratefully acknowledge the authors, originating and submitting laboratories of the sequences from GISAID’s EpiFlu™ Database on which this research is based. A table of the contributors is available in Dataset S1.

All submitters of data may be contacted directly via www.gisaid.org

## Supporting material

**Supplementary Table 1:** Linkage Disequilibrium linear model results. With bold face fonts we denote the populations characterized by p-values < 0.01 for the *a* parameter (the slope of the linear model).

| Population | *a* (p-value) | *b* (p-value) |
| --- | --- | --- |
| **All** | **-1.152e-07 (<2e-16)** | **3.944e-03 (<2e-16)** |
| **Europe** | **-1.905512e-07 (<2e-16)** | **6.344816e-03 (<2e-16)** |
| North America | -3.172727e-08 (0.113) | 5.861180e-03 (<2e-16) |
| South America | -2.704880e-07 (0.818) | 1.967903e-01 (<2e-16) |
| Africa | 3.3661e-06 (3.380589e-02) | 1.4271e-01 (1.4423e-11) |
| Oceania | 9.764460e-08 (0.648) | 2.821038e-02 (<2e-16) |
| **Asia** | **-1.494132e-06 (<2e-16)** | **3.608169e-02 (<2e-16)** |

**Supplementary Figure S1:**
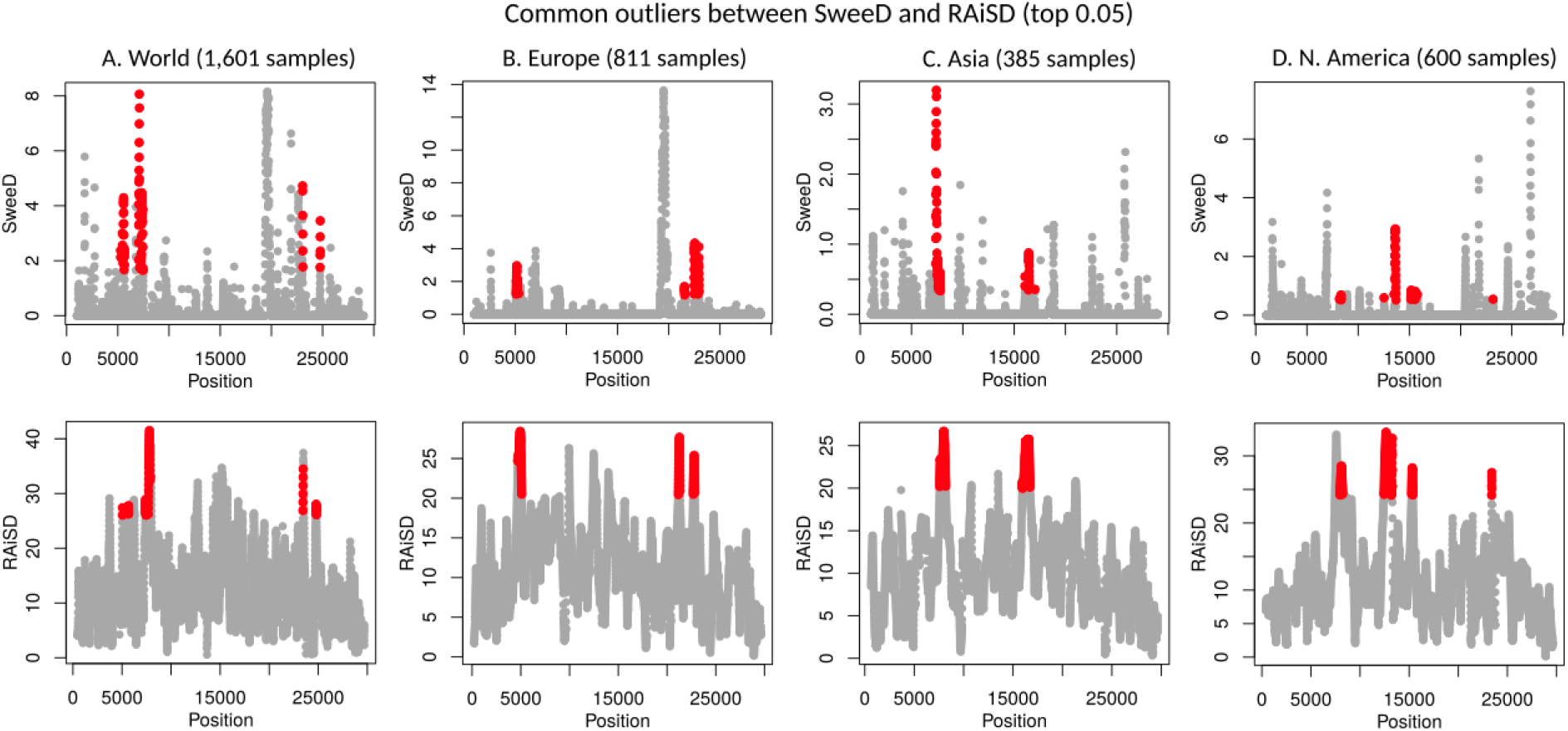
Common outliers between SweeD and RAiSD (top 0.05): A. World sample (1,601 genomes), B. Europe (811 genomes), C. Asia (385 genomes), and D. North America (600 genomes).

**Supplementary Figure S2:**
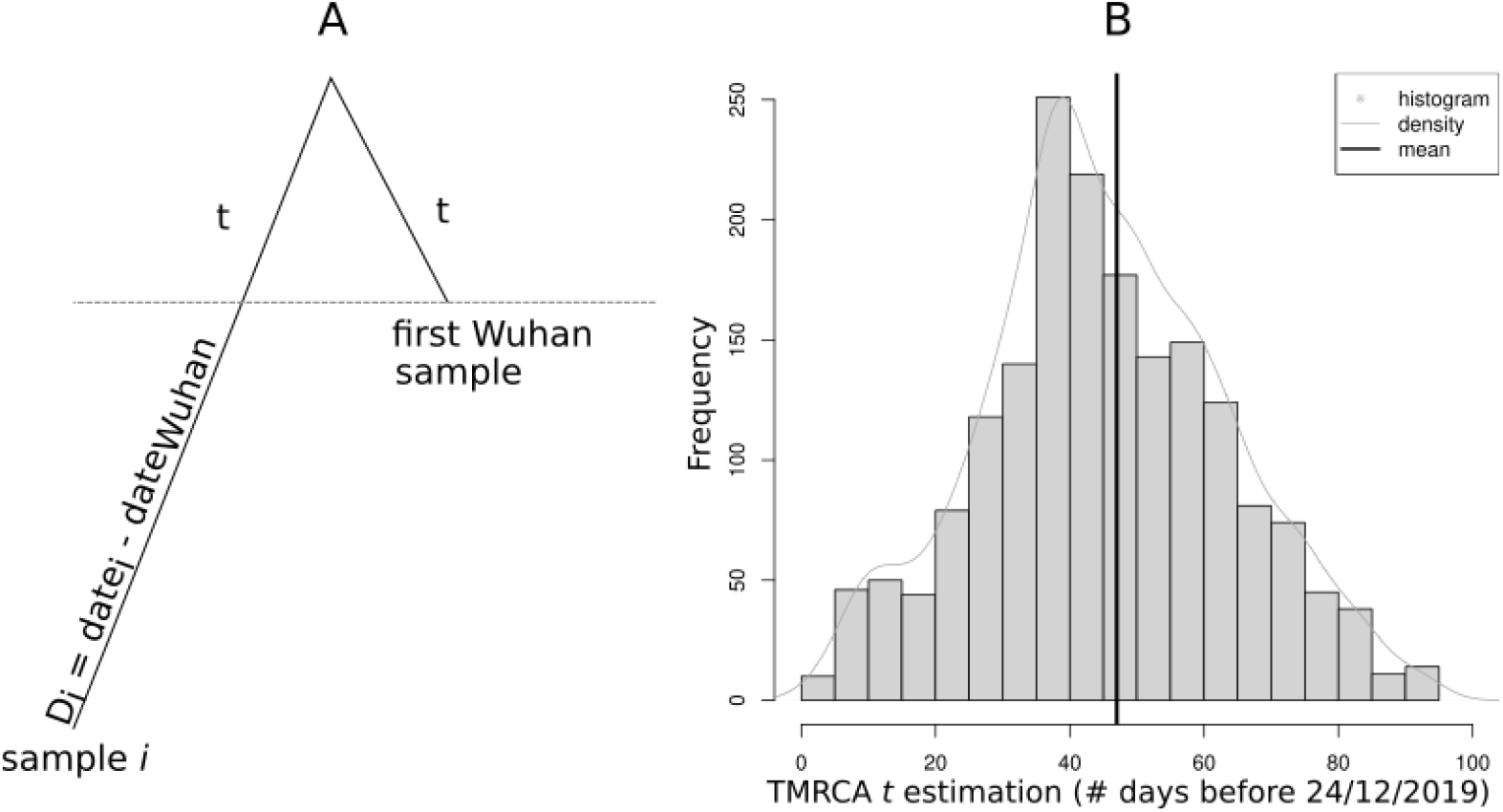
The parameter *t* denotes the time of the most recent common ancestor as estimated in units of days prior to 24/12/2019, the collection date of the first Wuhan sample that was reposited in GISAID.

**Supplementary Figure S3:**
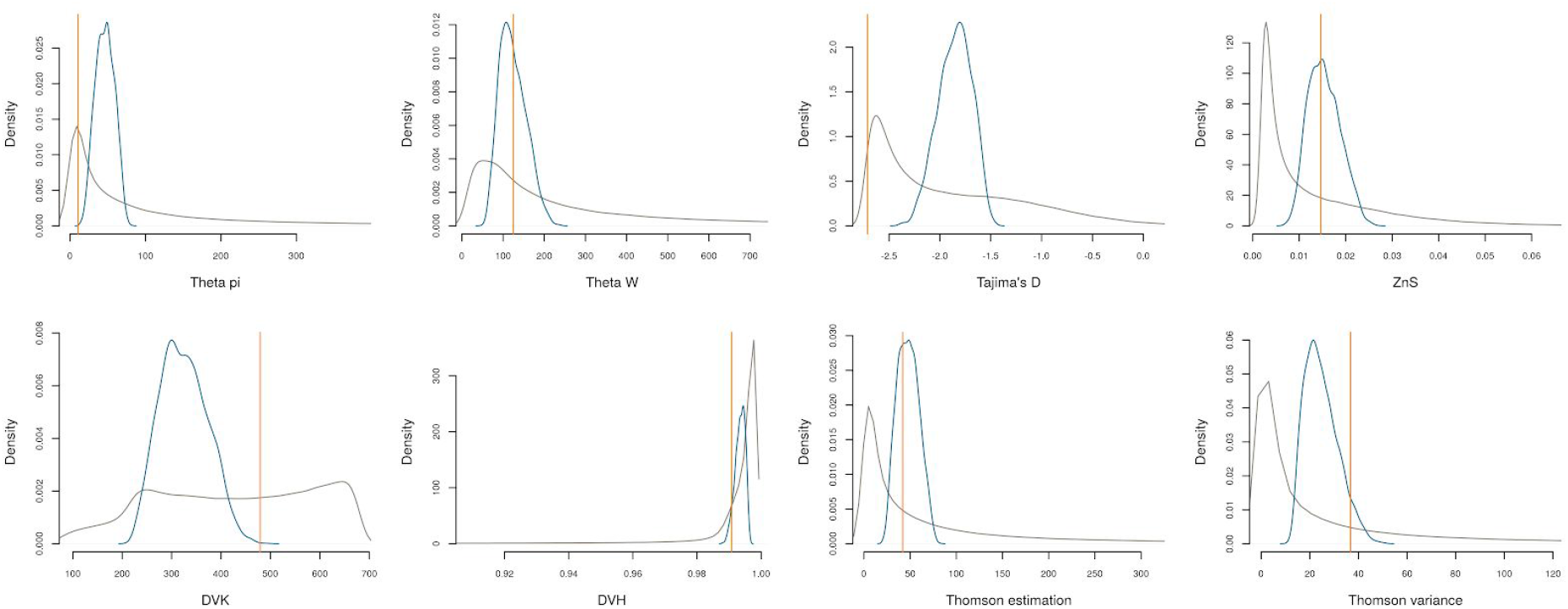
ABC parameter inference for the European population based on a set of 8 summary statistics. Orange line: Observed value of each summary statistic; Grey curve: density curve of the summary statistics calculated from the whole simulation space; Blue curve: density estimation of the ABC accepted summary statistics that were determined with a tolerance τ=0.005.

**Supplementary Figure S4:**
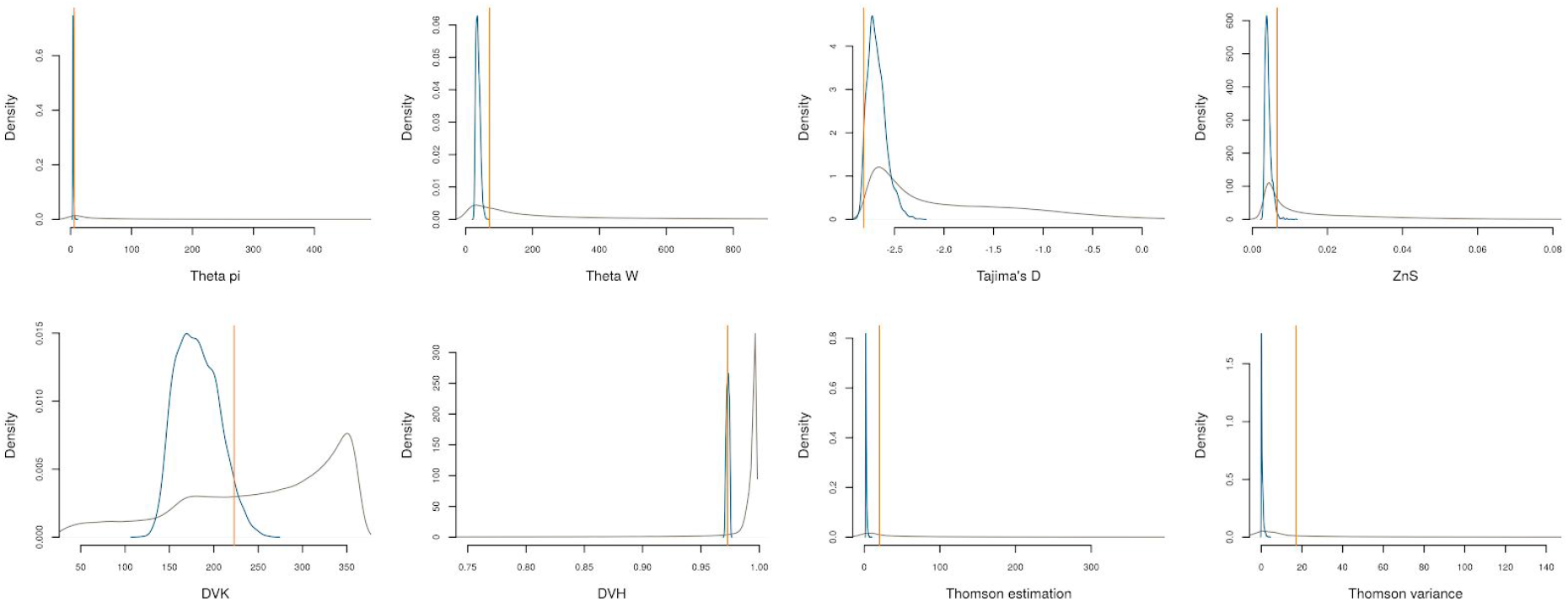
ABC parameter inference for the Asian population based on a set of 8 summary statistics. Orange line: Observed value of each summary statistic; Grey curve: density curve of the summary statistics calculated from the whole simulation space; Blue curve: density estimation of the ABC accepted summary statistics that were determined with a tolerance τ=0.005.

**Supplementary Figure S5:**
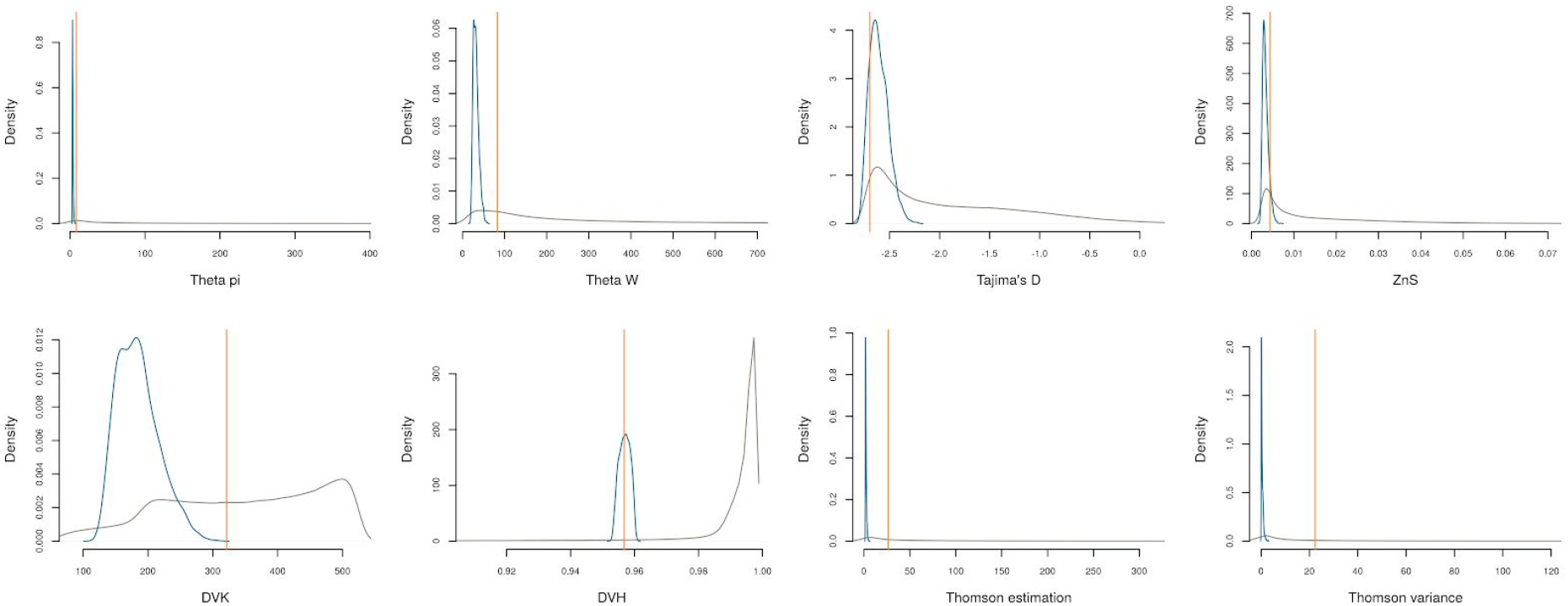
ABC parameter inference for the Northern American population based on a set of 8 summary statistics. Orange line: Observed value of each summary statistic; Grey curve: density curve of the summary statistics calculated from the whole simulation space; Blue curve: density estimation of the ABC accepted summary statistics that were determined with a tolerance τ=0.005.

**Supplementary Figure S6:**
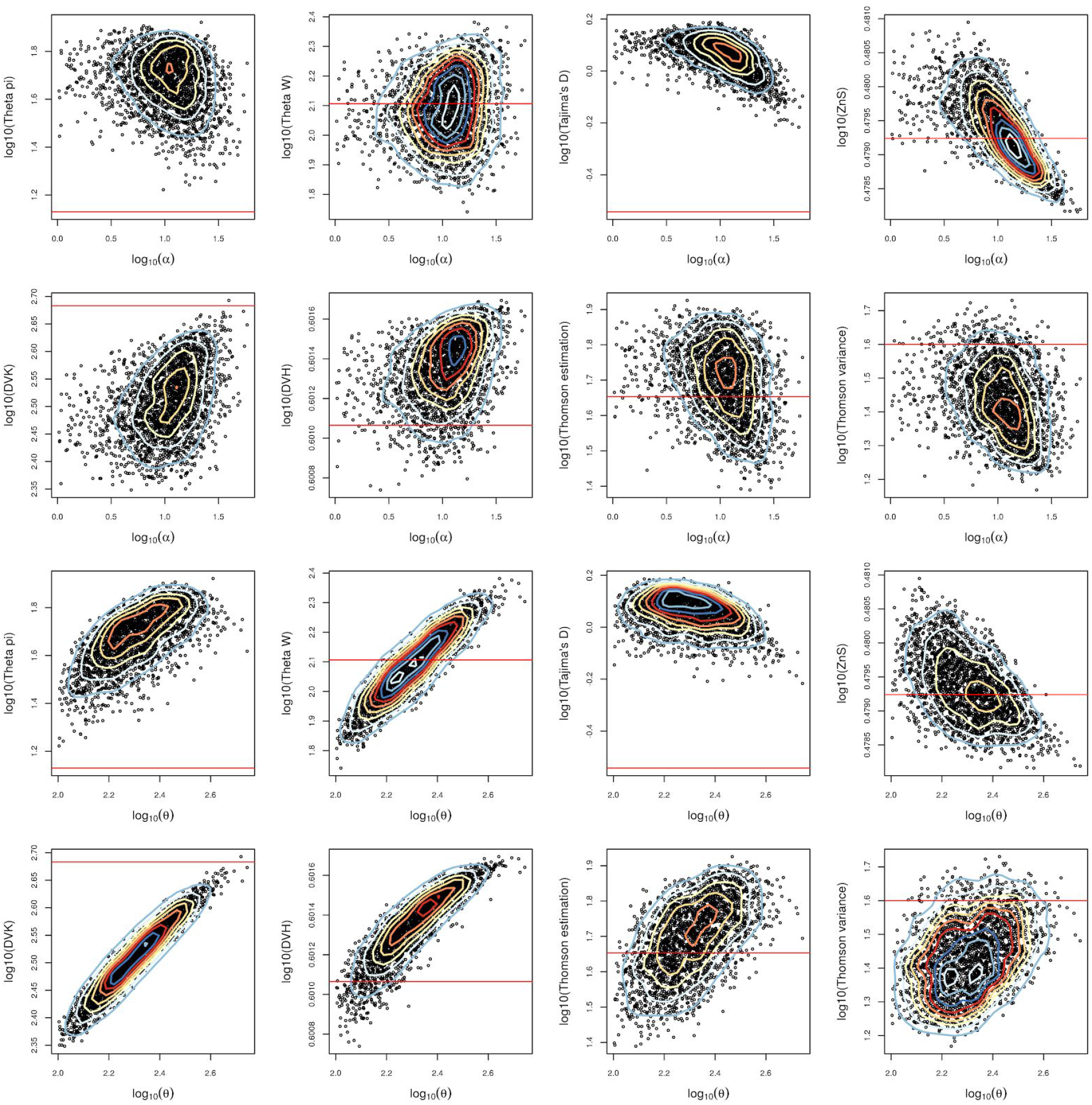
The space of the ABC accepted simulations of the European population. The contour lines on the scatterplot indicate the density estimation.

**Supplementary Figure S7:**
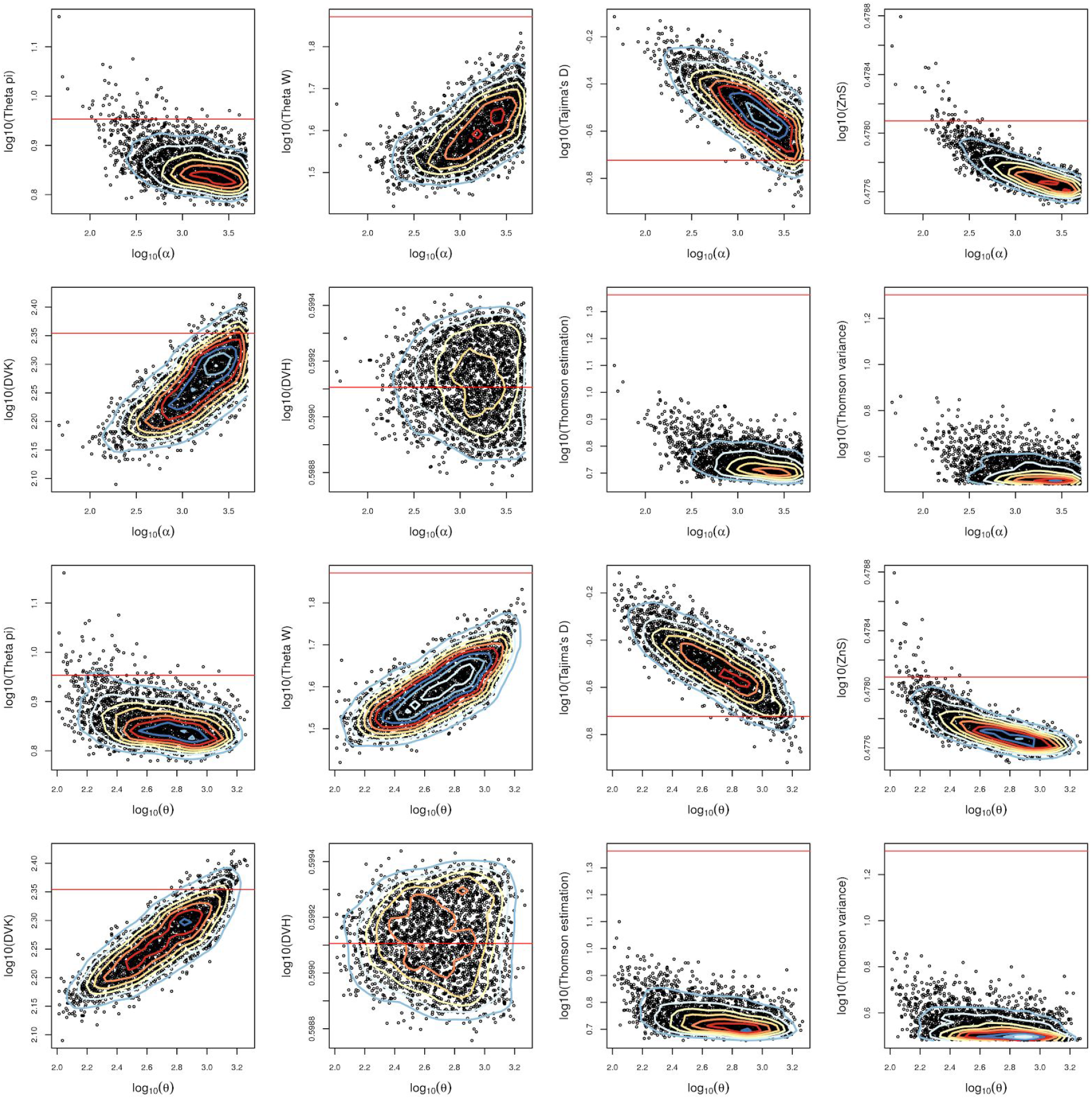
The space of the ABC accepted simulations of the Asian population. The contour lines on the scatterplot indicate the density estimation.

**Supplementary Figure S8:**
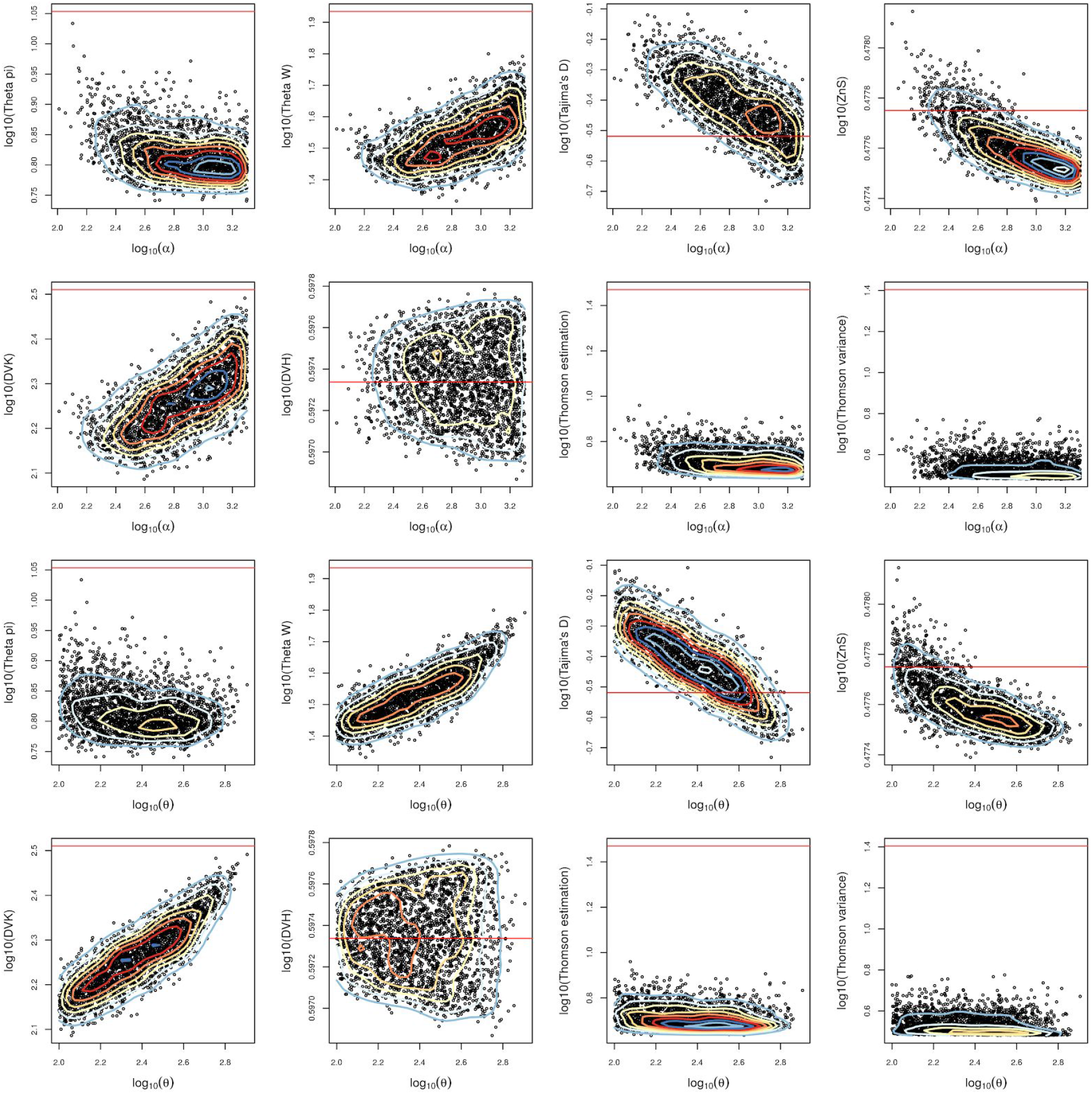
The space of the ABC accepted simulations of the Norhtern American population. The contour lines on the scatterplot indicate the density estimation.

**Supplementary Figure S9:**
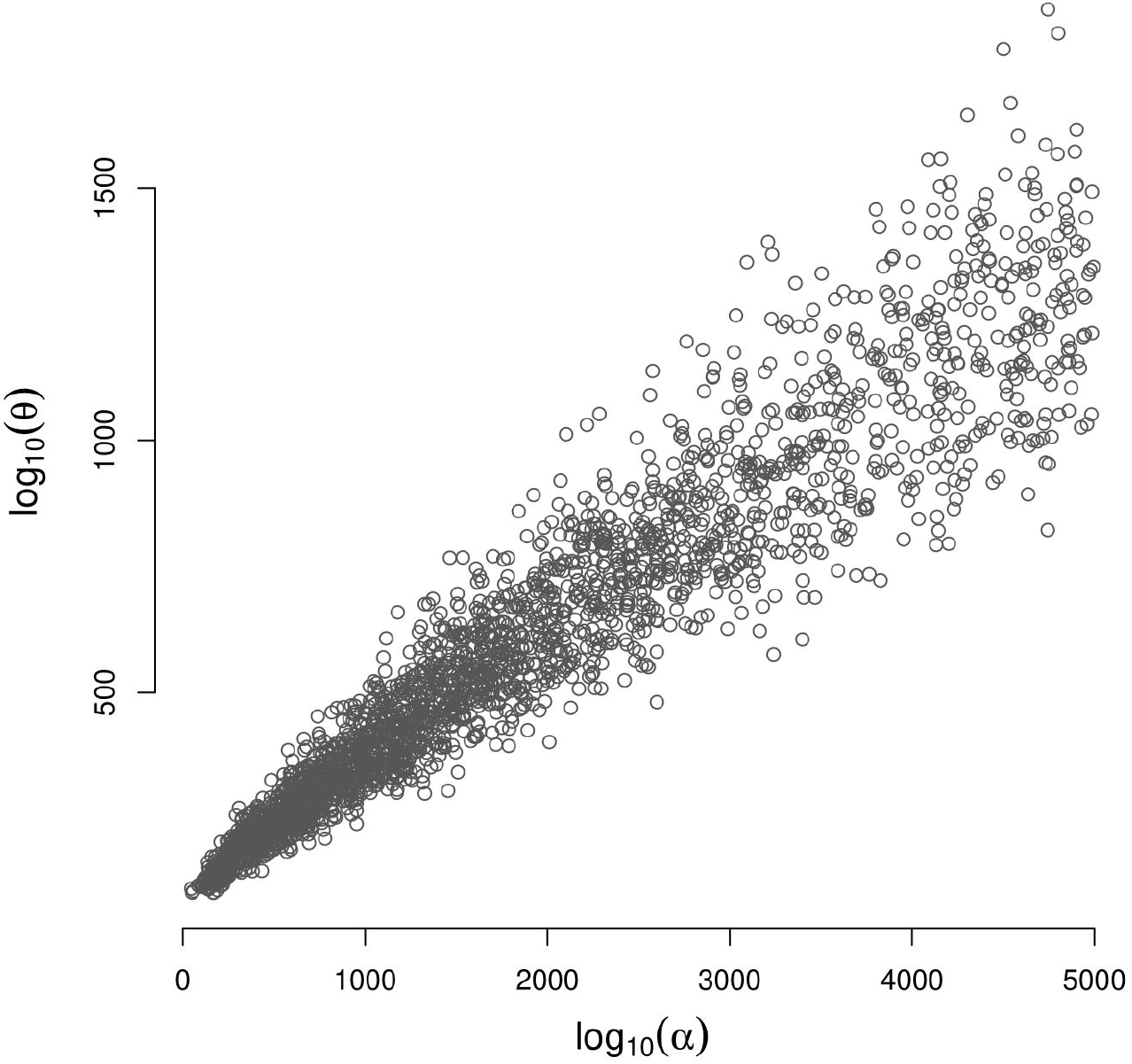
Correlation of Asia’s population α and θ parameters of the ABC accepted simulations (unadjusted values). The Pearson correlation coefficient is 0.95.

